# The *Ustilago maydis* transcription factor Nit2 regulates nitrate utilization during biotrophy and influences organic nitrogen metabolism in infected maize leaves under nitrogen limitation

**DOI:** 10.1101/2025.01.02.631094

**Authors:** Philipp L. Lopinski, Christin Schulz, Alicia Fischer, Nadine Reichl, Timo Engelsdorf, Nadja Braun, Lars M. Voll

**Author notes:** These authors contributed equally (Philipp L. Lopinski, Christin Schulz). **Corresponding author:** Lars M. Voll.

## Abstract

Little is known on how filamentous phytopathogens adapt to nitrogen limitation in the host plant. In previous work, we have shown that the transcription factor Nit2 plays a major role for the utilization of non-favored nitrogen sources like nitrate, minor amino acids or nucleobases in saprotrophic sporidia of the basidiomycete corn smut fungus *Ustilago maydis*.

Here, we employed *Δnit2* mutants in the natural FB1 x FB2 background to identify Nit2 regulated genes during biotrophy and investigated the impact of Nit2 on the physiology of leaf galls in nitrogen replete versus nitrogen limited host plants. RNA-Seq analysis of galls caused by the *U. maydis Δnit2* mutant and by the *U. maydis* wild type, revealed that about one third of the genes affected by Nit2 deletion during fungal biotrophy are involved in nitrogen metabolism and transport. Induction of the nitrate assimilation cluster was completely dependent on Nit2 and under nitrogen limitation, *Δnit2* leaf galls accumulated nitrate and showed reduced accumulation of the nitrogen-rich phloem transport amino acids asparagine and glutamine compared to wild type galls. In nitrogen replete conditions, only asparagine content was reduced in *Δnit2* leaf galls compared to wild type galls. Since total protein content in galls and pathogenicity were comparable between fungal genotypes in both nitrogen regimes, our findings demonstrate that nitrate utilization is dispensable for *Ustilago maydis* during biotrophy and can likely be compensated by increased utilization of abundant organic nitrogen sources, like asparagine, GABA and glutamine, which occurs in a partially Nit2-dependent fashion.

## Introduction

Fungal plant pathogens fuel the development of early infection structures by mobilization of lipid, protein and carbohydrate stores in spores (e.g. Both et al., 2005). For the ensuing establishment in the host tissue, biotrophic and hemibiotrophic pathogens have to gain access to nutrients from living host cells (as quite recently reviewed by Bezrutczyk et al., 2018). Photoassimilates like soluble sugars and amino acids have been identified as major organic nutrients for (hemi)biotrophic (fungal) leaf pathogens, but at the same time, represent important building blocks to fuel the host defence response. Thus, the competition for organic carbon and nitrogen represents a crucial battlefield in biotrophic plant-fungal interactions (as e.g. reviewed by Bolton, 2009; Fernandez et al., 2014; Bezrutczyk et al., 2018).

Several reports indicate that abundant organic nitrogen sources like amino acids are predominantly tapped for the nutrition of (hemi)biotrophs. Amino acid uptake transporters have been cloned from a range of fungal pathogens, e.g. seminal work for the rust fungus *Uromyces fabae* (Struck et al., 2002, Struck et al., 2004) or *C. fulvum* (Solomon and Oliver, 2002). GABA concentrations in the tomato apoplasm can reach up to 2 mM during *C. fulvum* infection (Solomon and Oliver, 2001), and circumstantial evidence suggests that it is the major organic nitrogen source of *C. fulvum* (Solomon and Oliver, 2002). Recently, the Arabidopsis amino acid transporter LHT1 was shown to be induced early during the host defense response against the hemibiotrophic bacterium *Pseudomonas syringae pv. syringae* in order to facilitate amino acid sequestration from the apoplast (Zhang et al., 2021), reflecting the aforementioned competition between host and parasite for apoplasmic amino acids. Highly interesting observations on the relevance of inorganic nitrate for nitrogen provision to phytopathogens derive from the comparative study of closely related oomycetes (Abrahamian et al., 2016; Ah-Fong et al., 2019). In particular, the nitrate assimilation cluster of *Phytophthora infestans* was strongly induced in nitrate-rich tomato leaves and deletion of the cluster resulted in complete loss of pathogenicity on leaves, while on nitrate-poor potato tubers, expression of the cluster was low and pathogenicity of the cluster mutants was mildly affected (Abrahamian et al., 2016).

In the corn smut fungus *Ustilago maydis*, biotrophy gets initiated during the dimorphic switch, in which two compatible haploid, saprotrophic sporidia are stimulated to mate by cues perceived from the host surface (Mendoza-Mendoza et al., 2006; Lanver et al., 2017a). Mating results in the formation of dikaryotic filamentous hyphae that develop appressoria-like structures, which can penetrate aerial organs of the host plant maize (reviewed by Lanver et al., 2017b). *U. maydis* is able to colonize meristematic tissue of maize leaves and reproductive organs with a network of inter- and intracellular hyphae that induce gall formation by stimulating cell divisions and hypertrophy in an organ-specific manner (Skibbe et al., 2010; Redkar et al., 2017). Due to easier handling and shorter cultivation times of host plants, nutrient acquisition and metabolic reprogramming have been studied most intensely in leaf galls. Interestingly, infection of leaf meristems by *U. maydis* prevents photoautotrophic development of maize leaves (Döhlemann et al., 2008), including the establishment of the C_4_ syndrome (Horst et al., 2008). Consequently, infected leaves do not develop into a physiological source organ but remain a sink for organic carbon and nitrogen until throughout gall development (Horst et al., 2008; Horst et al., 2010a). Using stable isotope labelling, Horst et al. (2010a) demonstrated that leaf galls outcompeted systemic sinks like young leaves for organic nitrogen and, in addition, stimulated amino acid export of systemic source leaves two-fold. For organic carbon acquisition, the *U. maydis* high-affinity H^+^/sucrose transporter Srt1 outcompetes host plant H^+^/sucrose transporters based on higher substrate affinity, which is pivotal for the establishment of biotrophy (Wahl et al., 2010).

Lanver et al. (2018) have elaborated a transcriptomic atlas for several stages of *U. maydis* biotrophy. Various transporters for inorganic and organic nitrogen, like the ammonia transporter *ump2* or the urea transporter *dur3-3*, were strongly induced during biotrophy, but deleting these genes did not affect virulence (Lanver et al., 2018). In addition, more than 60% of the encoded proton coupled co-transporters for hexoses and amino acids are induced during biotrophy of *U. maydis* (as by re-analysis of the data by Lanver et al., 2018), but functional data for their involvement in the provision of organic carbon and nitrogen to the pathogen are lacking to date, with one exception (Schuler et al., 2015). In contrast, the CRISPR/Cas9 induced mutual deletion of the oligopeptide transporter genes *opt2*, *opt3* and *opt4* diminished virulence (Lanver et al., 2018), indicating that these OPTs might contribute to the uptake of organic nitrogen during biotrophy.

In saprotrophic sporidia of *U. maydis*, the utilization of non-favoured nitrogen sources like minor amino acids, nucleobases and nitrate was shown to be predominantly controlled by the GATA Zn finger transcription factor Nit2 (Horst et al., 2012), indicating the presence of nitrogen catabolite repression (NCR) in sporidia. The involvement of *UmNit2* homologs in NCR of heterotrophic fungal model organisms like brewer’s yeast (*Saccharomyces cereviseae*), *Neurospora crassa* and *Aspergillus nidulans* is well described (reviewed in Wong et al., 2008). Several reports have demonstrated that Nit2 homologs are involved in the regulation of nitrogen utilization in plant pathogenic fungi (for reviews, see Divon and Fluhr, 2007; Fernandez et al., 2014). Intriguingly, most of the studied systems were hemibiotrophic ascomycetes. In *Colletotrichum lindemuthianum, Fusarium verticillioides* and *Fusarium oxysporum*, Nit2 homologs were required for full pathogenicity (Pellier et al., 2003, Divon et al., 2006; Kim and Woloshuk, 2008), while the loss of Nit2 homologs in *Magnaporthe grisea* or in *Cladosporium fulvum* had little or no effect on virulence (Froeliger and Carpenter, 1996; Perez-Garcia et al., 2001), but played a role for the expression of some effector proteins. For instance, the expression of the *C. fulvum* effector protein Avr9 was found to be under the control of the Nit2 homolog Nrf1 (Perez-Garcia et al., 2001) and five *M. grisea* pathogenicity factors were induced by nitrogen limitation *in vitro* (Talbot et al., 1993; van der Ackerveken et al., 1994; Donofrio et al., 2006). In the basidiomycete *U. maydis*, filamentation was delayed in *Δnit2* mutants in the solopathogenic strain SG200, resulting in reduced pathogenicity (Horst et al., 2012). Employing strains with an arabinose inducible *b-*cascade suggested that Nit2 acts downstream of the *b*-locus in the initiation of pathogenic growth during the dimorphic switch (Horst et al., 2012), indicating the integration of nitrogen signals into the regulation of pathogenicity of *U. maydis*.

In the present report, we aim at (i) identifying Nit2 regulated genes during biotrophy, as well as elucidating the role of Nit2 for (ii) nitrogen utilization and (iii) pathogenicity during biotrophy of *U. maydis* as well as (iv) its possible role for the adaptation to different nitrogen regimes and nitrogen limitation *in planta*. Investigating the involvement of Nit2 in the regulation of the dimorphic switch and the initiation of pathogenic growth is yet out of scope of the present manuscript.

## Materials and Methods

### Plant and fungal material and growth conditions

*Ustilago maydis* strains FB1Δ*nit2* and FB2Δ*nit2* were generated from FB1 and FB2 isolates (Banuett and Hershkowitz, 1994) by deleting the entire Nit2 open reading frame with the construct described in Horst et al. (2012). The *dur3-1,2,3* triple knockout strain was generously provided by Regine Kahmann (MPI for terrestrial microbiology, Marburg). Fungal strains were maintained on YEPS_light_ liquid medium and plates, while sporidia for plant infections were collected from potato dextrose plates (Tsukuda et al., 1988).

Maize *cv. Early Golden Bantam* was germinated in moist boxes at 28°C for 2 days, before seedlings of comparable size were transferred to P-type soil (Fruhstorfer Erde, Lauterbach, Germany) and plants were grown in phytochambers at a 14 h light (28°C, RH 50-60%) and 10 h dark (20°C, RH 60-70%) regime at a PFD of 350 µmol·m^-2^·s^-1^. We fertilized maize seedlings from 7-14 days post sowing with 1N regular Hoagland nutrient solution containing 16 mM nitrate and 1 mM ammonium (Table S1), or Hoagland nutrient solution with three-fold increased inorganic nitrogen concentration (3N) or Hoagland nutrient solution without any nitrogen source (-N), as indicated. Plant infection was conducted 7 days after transfer to soil as described (Gillissen et al., 1992; Horst et al., 2010a). To this end, equal volumes of compatible fungal sporidia (OD=1 each) were mixed and syringe-injected into the stems approx. 2 cm above ground, which resulted in the local infection of leaf 4 and sometimes also leaf 5. The same amount of water was injected into mock control plants.

### Isolation of total RNA and transcript analysis by RNA-Seq and qRT-PCR

Total RNA of gall material harvested at 8 dpi was extracted with the method of Chomzynski and Sacchi (1989) and RNA-Seq analysis was performed by Novogene (Cambridge, UK), while qRT-PCR validation of *U. maydis* regulated genes was performed as described by Horst et al. (2012), employing the primers described therein. Prior to their use in qRT-PCRs with samples from infected leaf material, all primer pairs were validated not to produce any products from non-infected leaf samples. During sample analysis by qRT-PCR, negative control samples from mock treated control leaves were always analyzed in parallel to rule out the contribution of plant derived amplificates.

### Isolation of genomc DNA and quantification of fungal DNA content by qPCR

Genomic DNA was extracted from medium sized galls at 8 dpi using the NucleoSpin Plant II Kit (Macherey-Nagel, Düren, Germany) according to the manufacturer’s instructions. Fungal and host DNA were quantified using the primer pairs for *UmPPI*, *UmRab7* and *ZmGAPDH as* described by Lanver et al. (2018).

### Quantification of protein, nitrate, ammonium and free amino acid contents

At 8 dpi, medium sized galls located on the middle of the blade of infected fourth leaves were excised and employed for steady state metabolite analysis along with samples taken from the same position and leaf number of mock treated control plants. Analysis of protein and nitrate were performed as described by Horst et al. (2010a). Extraction of leaf material and derivatization of ammonium and free amino acid with AQC (6-aminoquinolyl-N-hydroxysuccinimidyl carbamate) was also conducted according to Horst et al. (2010a).

HPLC separation of AQC derivatives was performed on a Macherey-Nagel (Düren, Germany) EC 150/3 NUCLEOSHELL RP 18plus (2.7 µm, 150mm length, 3.0 mm internal diameter) C_18_ column equipped with an EC UNIVERSAL RP guard column at a column temperature of 37 °C and a flow rate of 1.1 ml/min on a Agilent 1260 Infinity II Prime UHPLC system (Santa Clara, USA) by a ternary buffer system with buffer A (140 mM sodium acetate + 7 mM triethanolamine, pH 6.0), buffer B (99% acetonitrile) and buffer C (H_2_O_dd_). The gradient started with 98% A, 2% B, reaching 94% A and 6% B at 17 minutes, 88% A and 12% B at 20 minutes, 80% A and 20% B at 29 minutes, before the column was washed with 60% B and 40% C from 32 to 34 minutes, before the column was equilibrated to 98% A and 2% B until 38 minutes, which was kept for 2 minutes until the total run time of 40 minutes. Detection was performed with an Agilent G712B fluorescence detector at an excitation wavelength of 300 nm and an emission wavelength of 400 nm. For quantification, dilution series of the Pierce amino acid standard supplemented with GABA were analysed in the range of 1-200 pmol per injection.

### Statistical analysis

Statistical analysis of the data was performed with Graph Pad Prism (10.4.1) employing two-way ANOVAs with Fisher LSD post hoc tests.

## Results

### Establishment of a low N matrix for *U. maydis* biotrophy

To address the role of Nit2 for the adaptation to different nitrogen regimes and, ultimately, potential nitrogen limitation *in planta*, we aimed at generating a high N and low N environment for *U. maydis* during biotrophy in host leaves. Therefore, we modified the fertilization regime of the host plants and characterized the availability of free and protein bound amino acids as a proxy for overall availability of organic nitrogen, which we had already benchmarked in a previous study (Horst et al., 2010a). To this end, we fertilized maize seedlings from 7-14 days post sowing with 1N regular Hoagland nutrient solution containing 16 mM nitrate and 1 mM ammonium (as in Horst et al., 2010a, 2012), Hoagland nutrient solution with three-fold increased inorganic nitrogen concentration (3N) and Hoagland solution without any nitrogen source (-N). In nitrogen replete conditions (1N and 3N), the 4^th^ leaves showed significantly increased free amino acid contents compared to the 3^rd^ leaves (Figure 1), which reflects that the 4^th^ leaves are the first true C_4_ leaves. Moreover, there were no substantial differences between 1N and 3N regimes for both leaf positions, while in the -N regime, free and protein bound amino acid content was reduced two- and three-fold compared to 1N in 3^rd^ and 4^th^ leaves, respectively (Figure 1). Since the main objective of our study was to investigate the impact of nitrogen limitation on gall metabolism and gene regulation in *U. maydis Δnit2* mutants, we chose to further investigate galls on the 4^th^ leaves in -N and 1N, as the effect of nitrogen limitation on the biological matrix was most pronounced in this pair of conditions. Since there were comparatively minor differences in steady state nitrogen metabolism between 1N and 3N conditions, the 3N regime was discontinued in further experiments.

**Figure 1.**
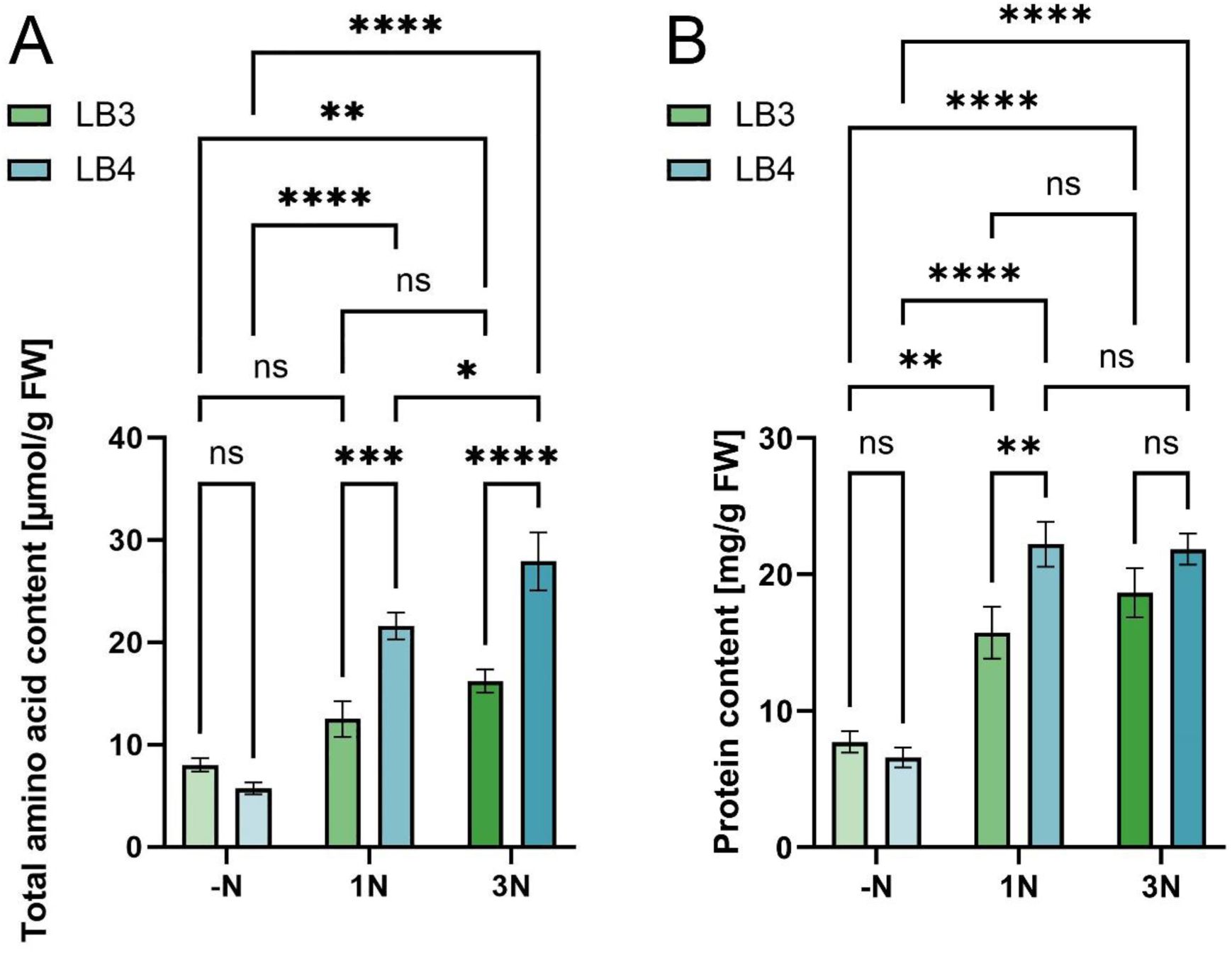
Contents of free and protein bound amino acids in maize leaves in the examined fertilization regimes. The contents of **(A)** total free amino acids and **(B)** protein content in leaf 3 (green bars) and leaf 4 (blue bars) of 14 days old maize plants that had been fertilized from day 7 with Hoagland nutrient solution with no nitrogen source (-N, left pairs of bars), 16 mM NO ^-^ and 1 mM NH ^+^ (1N, middle) and threefold increased inorganic nitrogen (3N, right). Data are means of four biological replicates (n=4) with the error bar representing the SE. Statistical analysis was performed with a two-way ANOVA and a Fisher LSD post hoc test (*P < 0.05; **P < 0.01; ***P < 0.001; ****P < 0.0001).

### The *Δnit2* mutant does not show reduced virulence in the wild type background

Since the pivotal aim of our study was to elucidate Nit2 regulated genes during late biotrophy, we decided not to use the previously described *Δnit2* mutant in the solopathogenic SG200 background (Horst et al., 2012), as defects during late stages of biotrophy were reported for the SG200 strain (Lanver et al., 2018). We therefore generated a full-length *nit2* knockout in the compatible FB1 and FB2 wild type strains, employing the *Δnit2* deletion construct described before (Horst et al., 2012). We confirmed the deletion of the entire Nit2 ORF by PCR (Figure S1) and the inability of the FB1Δ*nit2* and FB2Δ*nit2* mutants to utilize non-favoured nitrogen sources (Table S2). To assess, if the biological effect of Nit2 deletion on target gene regulation in FB1 and FB2 sporidia was quantitatively comparable to our previous observations for the SG200 background (Horst et al., 2012), we quantified transcript amounts of three selected Nit2 regulated genes by qRT-PCR in nitrogen deplete conditions (-NMM) versus nitrogen replete conditions with ammonia as a favourite nitrogen source (AMM), using oligonucleotide primers as described previously (Horst et al., 2012). The induction of the purine transporter *um01756*, the urea permease *dur3-3* (*um04577*) and the high-affinity ammonium transporter *ump2* (*um05889*) on -N compared to AMM was very similar between FB2 and SG200 as well as between FB2Δ*nit2* and SG200Δ*nit2* (compare Table S3 to Horst et al., 2012), while induction of *um01756* and *dur3-3* during N depletion were weaker in FB1 than in SG200 or FB2. The induction of *ump2* on –N versus AMM was comparable for all three control and all three Δ*nit2* strains, however.

Since we had observed delayed filamentation and reduced virulence of the SG200Δ*nit2* mutant (Horst et al., 2012), we next assessed the effect of *nit2* deletion on pathogenicity in the FB1 x FB2 wild type background,. Filamentation was also delayed 18 hours after mating FB1Δ*nit2* with FB2Δ*nit2* compared to the wild type FB1 x FB2 cross (Figure 2B), but symptoms at 8 dpi were similar between FB1Δ*nit2* x FB2Δ*nit2* and FB1 x FB2 on maize plants in both N regimes (Figure 2A), which was consistent between all six replicate experiments. Since this observation was in contrast to the results we obtained in the SG200 background (Horst et al., 2012), we further investigated whether fungal proliferation in the galls was similar between FB1Δ*nit2* x FB2Δ*nit2* and FB1 x FB2 by two complementary assays. Quantitation of fungal genomic DNA can be regarded as a proxy for the total number of fungal nuclei in the tissue (Figure 2C), while quantification of transcripts for fungal (*UmRab7*) relative to host (*ZmGAPDH*) housekeeping genes provides an estimate for transcriptionally active fungal nuclei (Figure 2D). Both on the DNA and mRNA level, no significant differences in colonization between mutant and wild type were detected in regular fertilized maize leaves (1N) at 8 dpi, with the mRNA data showing less variance than the genomic DNA data (Figure 2C and 2D, respectively).

**Figure 2.**
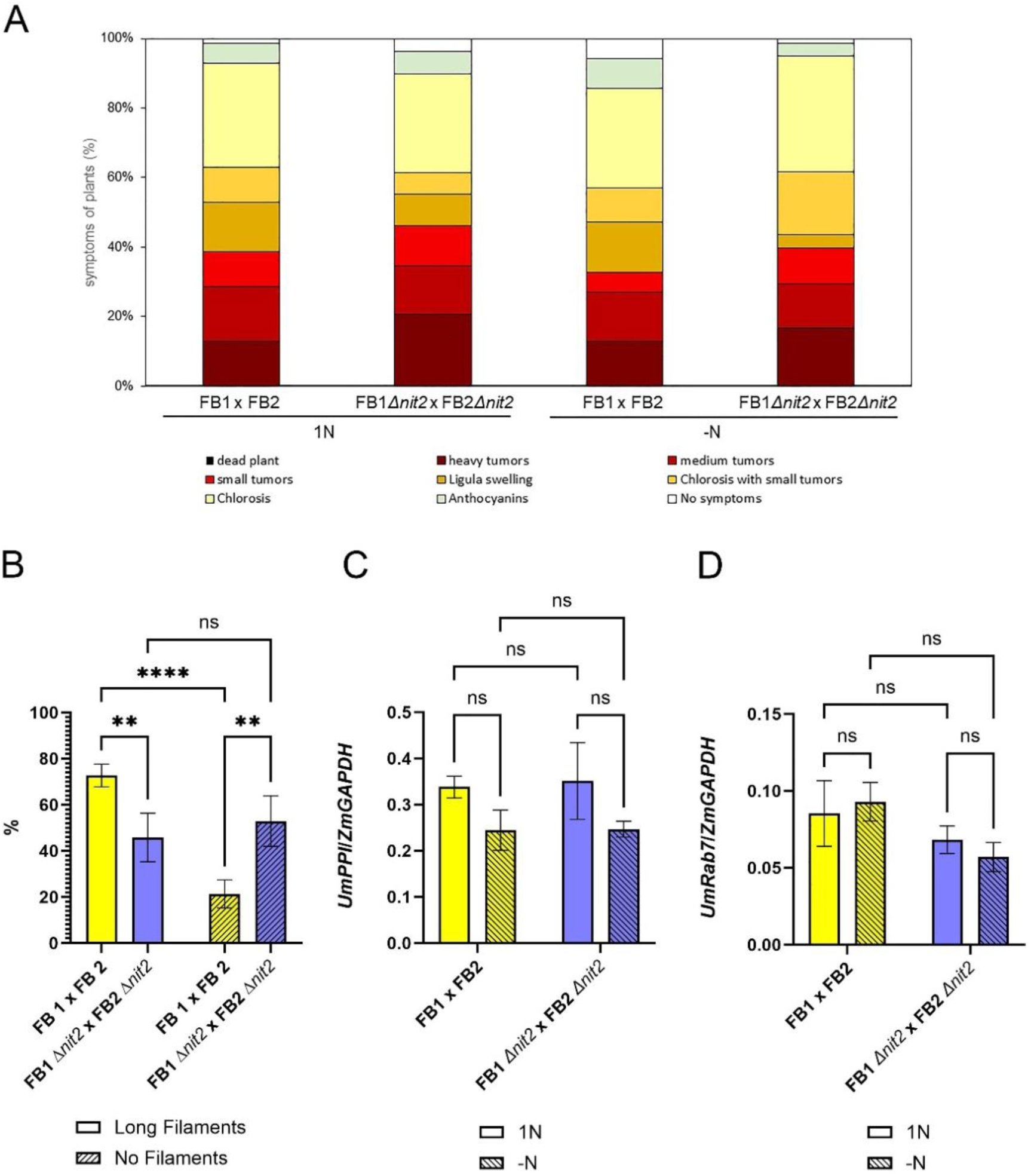
Pathogenic growth of FB1Δ*nit2* x FB2Δ*nit2* and FB1 x FB2 on maize leaves. **(A)** Disease index at 8 days post infection (dpi) with a mixture of the indicated sporidia at a titer of OD=1. N=70-78 with data from six independent experiment. **(B)** Filamentation on maize leaves 18 hours post inoculation with a mixture of the indicated sporidia at a titer of OD=1 was assessed by calcofluor white staining of fungal structures around the injection site. **(C)** Quantitation of fungal colonization in medium-sized galls on the fourth leaves of 14 days old plants at 8 dpi by qPCR on genomic DNA, employing fragments of the *UmPPI* and *ZmGAPDH* gene, data are means of six to eleven (n= 6-11) biological replicates with the error bar representing the SE. **(D)** Quantification of fungal colonization in medium-sized galls on the fourth leaves of 14 days old plants at 8 dpi by qRT-PCR for *U. maydis* and *Zea mays* housekeeping genes *UmRab7* and *ZmGAPDH*, respectively. Data are means of three to five biological replicates with the error bar representing the SE. For panels **(A)**, **(C)** and **(D)**, statistical analysis was performed with a two-way ANOVA and a Fisher LSD post hoc test (*P < 0.05; **P < 0.01; ***P < 0.001; ****P < 0.0001). Yellow bars: FB1 x FB2, blue bars: FB1Δ*nit2* x FB2Δ*nit2*, open bars: 1N, hatched bars: -N.

These observations indicate that despite delayed filamentation, the ability to colonize leaf tissue remains unaltered for a cross of FB1Δ*nit2* x FB2Δ*nit2* versus FB1 x FB2 at a saturating infection titer (OD = 1), which is an important prerequisite for an unbiased study of Nit2 regulated genes during biotrophy. It is highly interesting, but out of the scope of the present manuscript, to investigate the molecular basis why pathogenicity is compromised upon loss of Nit2 in the solopathogenic SG200, but not in the wild type strain, while filamentation is reduced to a similar extent in both backgrounds.

Even more interesting, the fertilization regime did not have a significant effect on disease development or host colonization (Figure 2A, 2C and 2D), which leads to the important conclusion that Nit2 is dispensable for pathogenicity on nitrogen limited host plants. However, it seems well possible that loss of Nit2 can be compensated, making it even more interesting to investigate Nit2 regulated genes during biotrophy.

### Identification of Nit2 regulated genes during biotrophy

To assess the potential difference in gene regulation by Nit2 between saprotrophic sporidia and biotrophic filaments *in planta*, we first investigated transcript accumulation for *um01756*, *dur3-3* (*um04577*) and *ump2* (*um05889*) representing those three Nit2 regulated genes that we had previously validated in sporidia of the FB1Δ*nit2* and FB2Δ*nit2* mutant (see Table S3). In order to investigate comparable tissue in a phase of well-established biotrophy, we harvested medium sized galls on the fourth leaves at 8 dpi for transcript analysis. Of these three analyzed genes, only *ump2* was induced in *U. maydis* infected leaves of –N treated maize plants compared to leaves from 1N fertilized maize plants, indicating a partially Nit2-dependent regulation in –N conditions *in planta* (Table 1). Furthermore, Nit2 seemed to be generally required for full induction of fungal *dur3-3* in galls: while normalized *dur3-3* transcript amounts significantly differed between FB1Δ*nit2* x FB2Δ*nit2* and FB1 x FB2 in 1N, a similar trend was observed in -N conditions (Table 1).

**Table 1.** Transcript accumulation of selected Nit2 regulated genes in sporidia during biotrophy at 8 dpi by qRT-PCR, with factor out 10^-2^. Transcript amounts were quantified relative to the *UmGAPDH* reference gene (Horst et al., 2012) and are means of 5-8 biological replicates ± SE. Statistical analysis was conducted with a two-way ANOVA and a Fisher LSD post hoc test. Treatments x genotype combinations that exhibit a significant difference to the other combinations with P < 0.05 are shown in bold.

| GeneID | Annotation | 1N |  | -N |  |
| --- | --- | --- | --- | --- | --- |
| | | FB1 x FB2 | FB1$\Delta nit2$ x FB1$\Delta nit2$ | FB1 x FB2 | FB1$\Delta nit2$ x FB1$\Delta nit2$ |
| | $\times 10^{-2}$ | | | | |
| *um01756* | purine transporter | $7.2 \pm 1.0$ | $5.3 \pm 0.4$ | $7.5 \pm 0.8$ | $6.3 \pm 0.6$ |
| *um04577* | urea permease *dur3* | $0.66 \pm 0.17$ | **$0.35 \pm 0.09$** | $0.73 \pm 0.14$ | $0.55 \pm 0.08$ |
| *um05889* | *ump2* | $7.2 \pm 1.2$ | $4.5 \pm 1.1$ | **$13.3 \pm 2.6$** | $5.6 \pm 0.8$ |

Taken together, we did not observe a strong Nit2-dependent regulation for all three tested genes in low N during biotrophy, suggesting a different set of Nit2 target genes *in planta*. Therefore, we performed RNA-Seq of *U. maydis* genes in medium sized leaf galls at 8 dpi on plants raised in 1N and –N conditions to screen for Nit2-regulated genes at a stage of well-established biotrophy (Table S4). In 1N conditions, we found 34 *U. maydis* genes showing a more than 2-fold reduction in transcript amount in FB1Δ*nit2* x FB2Δ*nit2* compared to FB1 x FB2 galls (Table S4), with 11 of these 34 genes, i.e. 32%, being involved in organic nitrogen metabolism, as based on annotation. In –N conditions, 44 *U. maydis* genes showed a more than 2-fold reduction in FB1Δ*nit2* x FB2Δ*nit2* compared to FB1 x FB2 galls (Table S4). Thirteen of these genes, approx. 30%, were associated with nitrogen metabolism. Among all genes, transcripts of *nit2* showed the strongest reduction of approx. 1000-fold in all 1N and –N samples from FB1Δ*nit2* x FB2Δ*nit2* galls compared to FB1 x FB2 Δ*nit2* galls, confirming a functional *nit2* knockout during biotrophy at 8 dpi. Notably, nitrite reductase *nir1* (*um11104*) transcripts were more than 30-fold diminished in Δ*nit2* galls in both N regimes, while nitrate reductase *nar1* (*um03847*) transcripts were almost 10-fold reduced in Δ*nit2* galls compared to FB1 x FB2 wild type galls in N deplete conditions (Table S4). In addition, the nitrate transporter *nrt* (*um11105*) showed an almost 30-fold decrease in Δ*nit2* galls compared to wild type galls in -N, and an 8-fold decrease in 1N conditions (Table S4). Therefore, we took a closer look on *nar1*, *nir1* and two other genes by qRT-PCR that showed the strongest Nit2-dependent regulation in both N regimes in an independent experiment (Figure 3).

**Figure 3.**
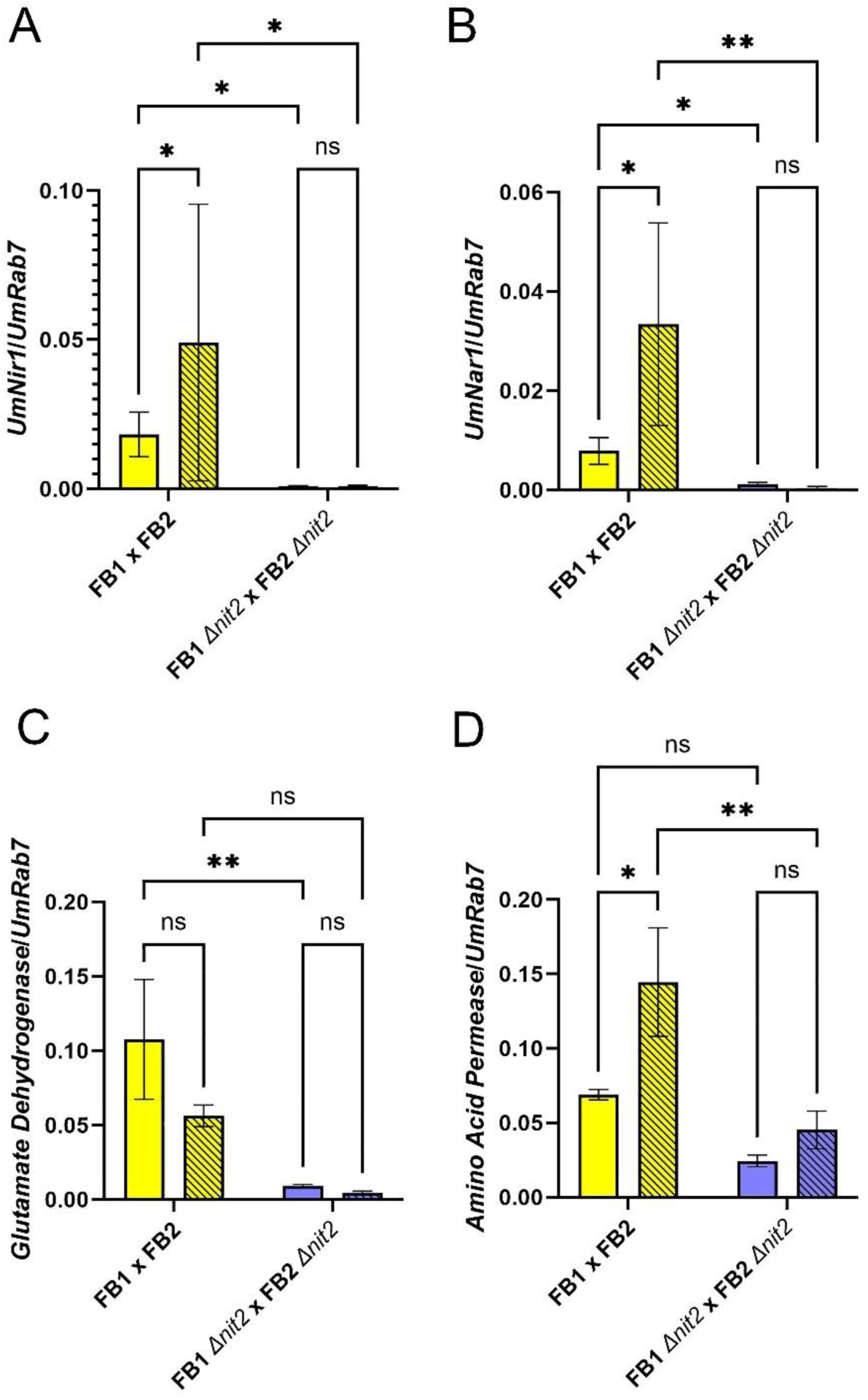
Nit2-dependent transcript accumulation of *U. maydis* genes in galls during biotrophy at 8 dpi. Medium galls of comparable size from the fourth leaf of 14 days old plants were harvested at 8 dpi and transcript amounts for the *U. maydis genes* **(A)** nitrite reductase *nir1* (*um11104*), **(B)** nitrate reductase *nar1* (*um03847*), **(C)** NADP glutamate dehydrogenase (*um02801*) and **(D)** an amino acid permease (*um00056*) were quantified in a qRT-PCR relative to the *Rab7* reference gene. Yellow bars: FB1 x FB2, blue bars: FB1Δ*nit2* x FB2Δ*nit2*, open bars: 1N, hatched bars: -N. Similar results were obtained when *PPI* was used as a reference gene (Figure S2). Values are means of 3-5 biological replicates ± SE. Statistical analysis was conducted with a two-way ANOVA and a Fisher LSD post hoc test (*P < 0.05; **P < 0.01; ***P < 0.001; ****P < 0.0001).

In galls at 8 dpi, the *U. maydis nir1* gene was induced 2.7-fold in the FB1 x FB2 wild type upon N depletion of the host (Figure 3A), while the *U. maydis nar1* gene was induced about 4-fold in the wild type (Figure 3B), confirming the nitrogen-dependent regulation of *nir1* and *nar1* during biotrophy of *U. maydis*. Furthermore, the results of the qRT-PCR confirmed the strong reduction in *nir1* and *nar1* transcript amounts in Δ*nit2* galls, especially in -N conditions (Figure 3A and 3B). Similar results for *nir1* and *nar1* were obtained when data were normalized to *PPI* instead of *Rab7* transcript amounts (Figure S2). Compared to the wild type, Δ*nit2* mutant galls showed an almost 20-fold reduction in *nir1* transcript and a 6-fold reduction in *nar1* transcript amount in regular fertilized plants (1N), while a more than 50-fold reduction and even a 93-fold reduction in *nir1* and *nar1* transcripts were evident in -N, respectively (Figure 3A and 3B), with the *nir1* transcript amounts being close to the detection limit in samples of Δ*nit2* galls. An N dependent regulation was absent for *nir1* and *nar1* in Δ*nit2* during biotrophy, indicating that nitrate utilization might generally be hampered in galls caused by Δ*nit2* mutants.

Transcript abundance for *U. maydis* NADP glutamate dehydrogenase (*um02801*) and the amino acid transporter *um00056* were also found to be diminished in Δ*nit2* compared to wild type galls at 8 dpi by qRT-PCR (Figure 3C and 3D). NADP glutamate dehydrogenase transcripts were about 10-fold reduced in Δ*nit2* galls in both N regimes (Figure 3C), while *um00056* transcripts accumulated 3-fold less in Δ*nit2* galls (Figure 3D). The amino acid permease *um00056* still showed a more than 2-fold induction in Δ*nit2* on -N compared to 1N grown host leaves, indicating the presence of other transcriptional regulators than *nit2*. In contrast, NADP glutamate dehydrogenase *um02801* rather appeared to be slightly transcriptionally repressed in N depleted host leaves compared to 1N fertilized leaves. While it can be assumed that the loss of Nit2 has substantial effects on nitrate utilization of *U. maydis* during biotrophy, the effect on the utilization of minor amino acids seems less apparent. Nevertheless, GO term analysis revealed that the categories ‘transport’ and ‘transmembrane transport’ were enriched among the *U. maydis* transcripts downregulated in Δ*nit2* compared to wild type galls in both N regimes (Figure S3). We therefore analyzed relative transcript abundance of all annotated amino acid transporters in the RNA-Seq datasets for wild type and Δ*nit2* in 1N and -N grown hosts (Figure S4). In the *U. maydis* wild type at 8 dpi, transcripts for *um00056* were the fourth and the fifth most abundant in 1N and -N, respectively, representing about 5% of all transcripts for amino acid transporters. For both wild type and Δ*nit2*, the top 5 candidates represented about half of all transcripts for amino acid transporter genes, with *um02549* and *um06012* alone amounting to 25% of the total. In -N, however, this pattern remained largely the same for Δ*nit2*, while for the wild type, the share of *um02549* and *um06012* increased to almost 50% (Figure S4), suggesting potential differences in amino acid uptake and metabolism in Δ*nit2* and wild type galls.

### Analysis of steady state amino acid contents in wild type and Δ*nit2 galls*

Steady state contents of free Asn, Gln and Glu (involved in N assimilation), GABA and Pro (derived from Glu) and the photorespiratory intermediate Ser were substantially increased in galls formed by both fungal genotypes on nitrogen replete maize plants at 8 dpi (Figure 4C - 4J), while contents of the C_4_ marker amino acid Ala were reduced by 50% in galls compared to mock control leaves on 1N fertilized plants (Figure 4E), similar to what was previously described in Horst et al. (2010a) for galls formed by SG200. In 1N conditions, only Asn contents were diminished in Δ*nit2* compared to wild type galls, while the steady state contents of all other amino acids did not differ between galls caused by the two genotypes in 1N.

**Figure 4.**
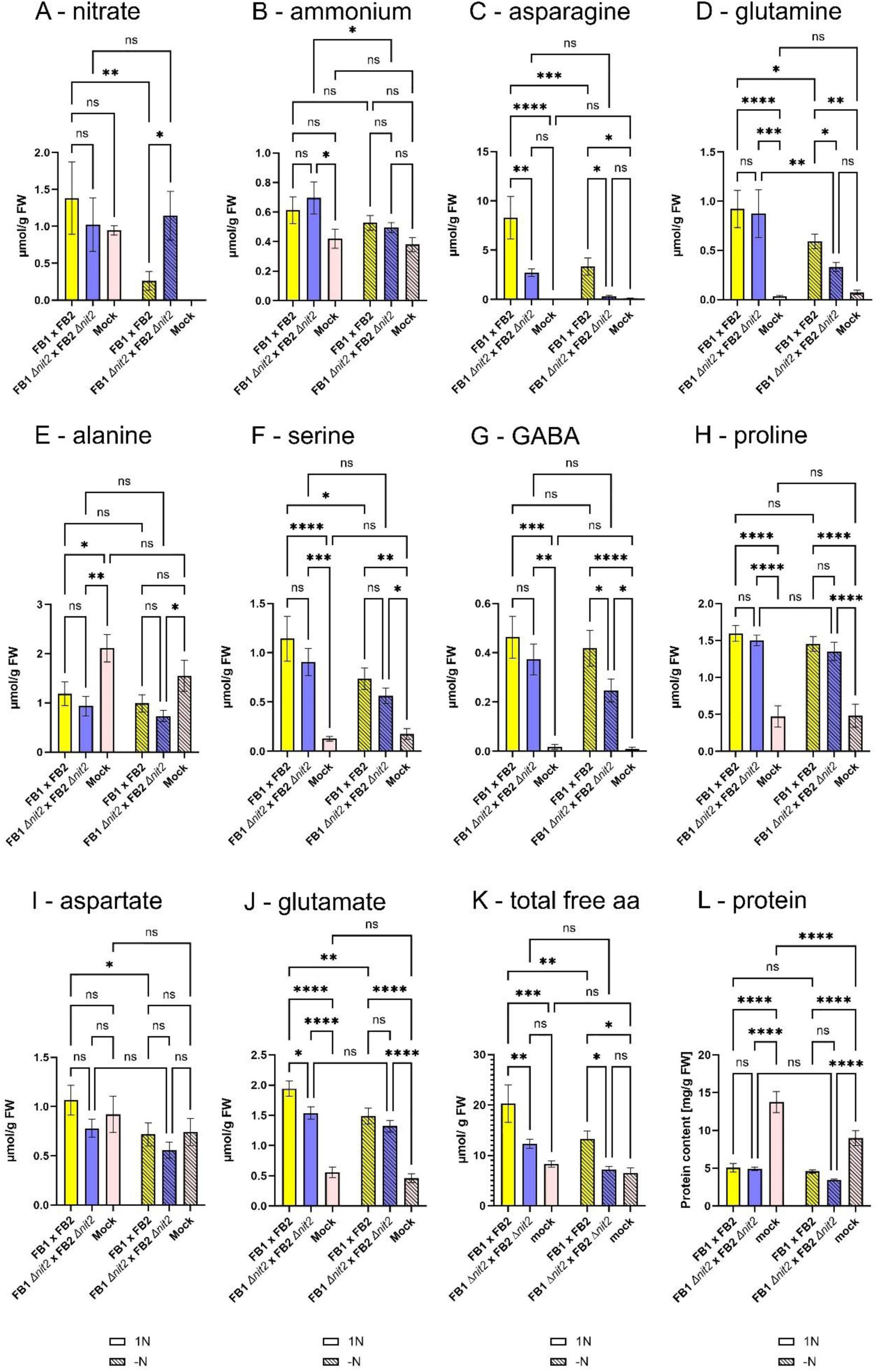
Steady state contents of major inorganic nitrogen sources as well as free and bound amino acids in medium sized galls at 8 dpi. **(A)** nitrate, **(B)** ammonium, **(C)** asparagine, **(D)** glutamine, **(E)** alanine, **(F)** serine, **(G)** GABA, **(H)** proline, **(I)** aspartate, **(J)** glutamate, **(K)** total free amino acid and **(L)** protein content. Medium galls of comparable size from the fourth leaf of 14 days old plants were harvested at 8 dpi. Yellow bars: FB1 x FB2, blue bars: FB1Δ*nit2* x FB2Δ*nit2*, pink bars: non-infected control leaves, open bars: 1N, hatched bars: -N. Values are means of 6-7 biological replicates ± SE for 1N, 8-11 biological replicates ± SE for –N and 4-6 biological replicates for healthy control leaves. Data from one representative out of three independent experiments are shown. Statistical analysis was conducted with a two-way ANOVA and a Fisher LSD post hoc test (*P < 0.05; **P < 0.01; ***P < 0.001; ****P < 0.0001).

In –N conditions, the total free amino acid content in mock control plants was 22% diminished compared to 1N mock controls (Figure 4K). In contrast, the contents of the major free amino acids Asn, Gln, Glu, GABA, Pro and Ser as well as total free amino acid content were increased in wild type galls compared to mock control leaves in -N, albeit to a lesser extent than in 1N (Figure 4), indicating that a strong flux of organic nitrogen is maintained into galls in N limited conditions. While the contents of most free amino acids were similar between wild type and Δ*nit2* galls, contents of Asn, Gln and GABA were decreased in Δ*nit2* compared to wild type galls in –N conditions, suggesting altered amino acid metabolism in Δ*nit2* galls. In turn, nitrate accumulated fourfold in Δ*nit2* compared to wild type galls in nitrogen deplete conditions. Furthermore, nitrate was not detectable in mock control leaves in –N (Figure 4A), indicating that nitrate is allocated to galls in –N, but cannot be utilized efficiently in Δ*nit2* galls. For comparison, nitrate contents were similar in Δ*nit2* galls, wild type galls and mock control leaves of nitrogen replete plants (Figure 4A). Free ammonium was slightly increased in galls compared to mock control leaves in both N regimes, with no difference between fungal genotypes (Figure 4B).

The observed free amino acid contents point towards confined differences in organic nitrogen metabolism between Δ*nit2* and wild type galls. To examine, if this has an impact on the capacity for protein synthesis, i.e. the pool size of protein bound amino acids, protein contents were determined (Figure 4L). Total leaf protein content was 25% reduced in –N compared to 1N grown mock treated fourth leaves, likely being a result of reduced nitrogen availability. Total leaf protein content was further reduced in galls, but did not differ between the N regimes and fungal genotypes (Figure 4L), indicating no substantial limitation in the availability of free amino acids for protein biosynthesis in galls of FB1Δ*nit2* x FB2Δ*nit2*.

## Discussion

### Disentangling the role of Nit2 in the control of pathogenicity and nitrate utilization during biotrophy

For the utilization of nitrate, a cluster comprising three genes, the nitrate transporter *nrt*, nitrate reductase *nar1*, and nitrite reductase *nir1*, is required in *Ustilago maydis* (McCann and Snetselar, 2008; Khanal et al., 2021). In sporidia, all three genes of the cluster, and hence nitrate utilization, were found to be under the predominant control of Nit2 in conditions when NCR was released, i.e. on nitrate minimal medium or upon nitrogen depletion (Horst et al., 2012). Studies on the role of nitrate utilization for *U. maydis* biotrophy have not only been hampered by the fact that Δ*nit2* mutants exhibited delayed and reduced filamentation in the solopathogenic SG200 background (Horst et al., 2012), but also because defects in any of the three structural genes of the nitrate utilization cluster resulted in apathogenic strains or strains with strongly reduced virulence (e.g. Khanal et al., 2021, own unpublished results). In other filamentous phytopathogens, defects in nitrate utilization did not affect pathogenicity in hemibiotrophic ascomycete fungi like *C. fulvum*, *Magnaporthe grisea* and *Stagonopora nodorum* (Cutler et al., 1988; Lau and Hamer, 1996; Talbot, 1990). Deleting the nitrate assimilation cluster in the oomycete *Phytophthora infestans* completely abrogated virulence in nitrate-rich tomato leaves, while virulence on nitrate-poor potato tubers was merely affected (Abrahamian et al., 2016). In most of the studied systems, nitrate utilization was dispensable for full virulence.

In the FB1 x FB2 background reported here, disruption of *Nit2* also lead to a strong reduction in filamentation as in the solopathogenic strain SG200, but did not reduce pathogenicity in the further course of the interaction. Given the high infection titer of OD=1, it is not surprising that a quantitative, minor delay in filamentation does not result in diminished colonization, since the number of timely successful penetration attempts will still be sufficient to allow for efficient colonization in the absence of further defects. This observation further suggests two things. First, this indicates that the artificial regulation of *b* in the solopathogenic SG200 strain might exert side effects on either penetration of post penetration growth. Interestingly, nitrogen limitation in haploid sporidia lead to ectopic activation of bE and bW in the 1/2 (*a1b1*, as FB1) but not in the 2/9 (*a2b2*, as FB2) background (Khanal et al., 2021), indicating different recruitment of Nit2 during the induction of filamentation by the two *b* alleles. Second, the observed loss of pathogenicity for mutants in nitrate metabolism might probably solely be due to a compromised early infection phase (as reported by Khanal et al., 2021). It will be highly interesting to resolve, how the interaction of *Nit2* and *b* proceeds on the molecular level, but this is clearly out of scope of the present manuscript, which is meant to describe the role of Nit2 for the regulation of nitrogen utilization during biotrophy.

To this end, we have observed that *nar1* and *nir1* are controlled by Nit2 during the biotrophic phase of *U. maydis*, both in regularly fertilized host leaves (1N) and in nitrogen depleted host leaves (-N), with consistent data obtained in independent experiments by RNA-Seq and qRT-PCR. Both, *nar1* and *nir1* transcripts were close to or at the detection limit in Δ*nit2* galls, indicating that nitrate utilization during biotrophy depends on Nit2. The degree of control exerted by Nit2 on *nar1* and *nir1* during biotrophy is much stronger in comparison to sporidia (Horst et al., 2012). It appears tempting to speculate that such a strong transcriptional control of nitrate utilization by Nit2 might have implications for the adaptation to nitrogen availability, but Nit2 was only 1.6-fold induced in wild type galls in -N compared to 1N conditions on the transcriptional level (Table S4), suggesting that either posttranslational regulation of *Nit2* prevails, or that the compared N regimes do not lead to a strong difference in the regulation of *Nit2*. In order to test if *Nit2* induction can be further increased during biotrophy, nitrogen limitation in the host plant could be exacerbated by applying -N conditions to plants cultivated on sandy or peaty substrates or by low light.

### Nitrate utilization is dispensable for pathogenicity during biotrophy

In order to unlock viable approaches for disease protection, crucial processes in the nutritional strategies of individual pathogens need to be identified *in planta*. Biotrophic phytopathogens rely on the active provision of nutrients, organic carbon and nitrogen sources by the host tissue, for which they actively reprogram host metabolism (for quite recent reviews see Fernandez et al., 2014; Bezrutczyk et al., 2018). Restricting supply of essential carbon building blocks from the host can mean a valuable addition to broad spectrum and durable resistance, as exemplified by the wheat *Lr67* gene (Moore et al., 2015) and the rice *xa13* gene (Chu et al., 2006; Yang et al., 2006; for a compilation see e.g. Bezrutczyk et al. 2018).

In the vast majority of analyzed pathosystems, fungal (hemi)biotrophs divert abundant organic nitrogen sources like amino acids as their preferred nitrogen source (see introduction), while the role of nitrate for pathogen nutrition is less well investigated. According to Solomon and Oliver (2001), the tomato apoplasm contains 4-5 mM nitrate, which makes it much more abundant than any proteinogenic amino acid, which are ranging between 0.1 - 0.7 mM. GABA concentrations in the tomato apoplasm can reach up to 2 mM during *C. fulvum* infection (Solomon and Oliver, 2001), and besides GABA being a potent ROS quencher, circumstantial evidence suggests that it is the major organic nitrogen source of *C. fulvum* (Solomon and Oliver, 2002). However, the contribution of nitrate to *C. fulvum* nitrogen acquisition has not been investigated. In the oomycete *Phytophthora infestans*, the nitrate assimilation cluster was strongly induced in nitrate-rich tomato leaves, but not in nitrate poor tubers (Abrahamian et al., 2016). Consequently, deletion of the nitrate assimilation cluster resulted in complete loss of pathogenicity on leaves, while pathogenicity was only mildly affected in tubers (Abrahamian et al., 2016). A subsequent study indicated that transcript accumulation of nitrate reductase and a homolog of the Nit2 repressor from filamentous fungi, NMR, were inversely correlated (Ah-Fong et al., 2019), indicating the control of nitrate utilization during pathogenic growth of *P. infestans* by NCR. In addition, metabolomic analysis of infected potato tubers indicated that several pathways for minor amino acid biosynthesis were strongly induced during the biotrophic phase of *P. infestans* (Ah-Fong et al., 2019), while 20% of all amino acid transporters were strongly expressed in tubers (Abrahamian et al., 2016), indicating that *P. infestans* might preferentially utilize free amino acids as nitrogen source in tuber tissue. Taken together, this indicates that *P. infestans* seems capable to adapt itself to utilize nitrate as predominant N source in response to the biological matrix.

In the maize apoplasm, nitrate and total amino acids concentrations were found to be similar at around 1.3 mM, while the leaf content of free amino acids was determined to be 5 times higher, i.e. 9.8 µmol/ g FW, than the leaf nitrate content of 1.8 µmol/ g FW (Lohaus et al., 2000). In the present study, we observed a free amino acid content of 8.3 µmol/ g FW and a leaf nitrate content of around 1.0 µmol/ g FW in mock control leaves in the 1N regime, which is in good accordance with the data by Lohaus et al. (2000). Therefore, we assume that the concentrations of amino acids and nitrate in the leaf apoplasm can also be deemed comparable between these two studies. While nitrate content was similar in galls and mock leaves in 1N conditions, nitrate could not be detected in mock leaves of nitrogen depleted plants, indicating a strong effect of the -N regime on foliar nitrate. Interestingly, nitrate was still detectable in galls under -N conditions, albeit nitrate contents in wild type galls were seven-fold reduced compared to 1N. In contrast, the nitrate content of Δ*nit2* galls remained similar in both N regimes. This suggests that (i) *U. maydis* colonization in galls generates a sink for nitrate and that (ii) this nitrate cannot be utilized by Δ*nit2* galls. Combined transcriptome and NR activity analysis by our group clearly indicated that nitrate assimilation by host cells is strongly reduced in galls compared to mock control leaves at 8 dpi (Horst et al., 2010a), which is probably caused by an overall suppression of photosynthetic development (Doehlemann et al., 2008) and associated pathways for organic carbon and nitrogen assimilation (Horst et al., 2008; Horst et al., 2010a). A parallel RNA-Seq analysis of maize genes did not show differential regulation of the host nitrate reductase gene *Nar1S* between Δ*nit2* galls and wild type galls in 1N or -N (not shown). Since the Δ*nit2* strain does not contribute any NR activity, we can conclude that diminished nitrate assimilation in host leaves causes nitrate accumulation in Δ*nit2* galls in -N conditions. As we observed five-fold less nitrate accumulation in galls caused by the wild type compared to Δ*nit2* galls, we conclude that the *U. maydis* wild type assimilates nitrate during biotrophy. And because fungal colonization did not differ between the two strains in -N, nitrate assimilation by *U. maydis* must be dispensable for virulence and biotrophy in nitrogen limited host plants.

### Nit2 and its downstream genes are dispensable for pathogenicity during biotrophy

While we found that nitrate assimilation by *nar1* and *nir1* was entirely under the control of Nit2 during biotrophy, we also observed that more than 30% of all differentially regulated genes between Δ*nit2* and wild type galls were associated with organic nitrogen metabolism (see Table S4). Most of these genes differed between 2 and 8-fold between the fungal genotypes, indicating that they were only partially controlled by Nit2 during biotrophy. Transcripts for three amino acid transporter genes, the genes for the GABA permease UGA4 related transporter *um03522*, and the two general amino acid transporters *um00056* and *um00343* were less abundant in Δ*nit2* galls compared to wild type galls in -N conditions. In contrast, only *um00056* and the general amino acid permease *um06012* were less expressed in Δ*nit2* galls in 1N conditions, indicating that Nit2 might play a role in remodeling amino acid uptake specificity of biotrophic hyphae in response to N availability. In fact, these adjustments in the transportome might prevent an effect on pathogenicity, as host colonization was found to be comparable between FB1 x FB2 and FB1Δ*nit2* x FB2Δ*nit2* as well as between nitrogen regimes at 8 dpi. Therefore, one or a group of these differentially regulated amino acid transporters might have an effect on pathogenicity when knocked out.

Other organic nitrogen sources might also contribute to nitrogen provision to *U. maydis*. The three putative oligopeptide transporters *opt2*, *opt3* and *opt4* were strongly induced during biotrophy and simultaneous CRISPR-Cas9 mediated knockout of all three homologs specifically affected pathogenicity in the late biotrophic phase (Lanver et al., 2018). OPTs can have a broad, but distinct substrate specificity, ranging from dipeptides to glutathione, phytochelatins and oligopeptides (e.g. Osawa et al., 2006; Ito et al., 2013). Pending the functional characterization of these three transporters *in vivo*, we cannot rule out that OPTs have additional vital functions than providing organic nitrogen during *U. maydis* biotrophy.

The same study by Lanver et al. (2018) revealed that, in contrast, knockout of all three *dur3* urea transporter homologs did not affect compatibility in nitrogen replete host plants. While two of the three encoded *dur3* urea permeases, *dur3-2* and *dur3-3*, were even partially controlled by Nit2 in 1N conditions, they were not found to be differentially regulated between the strains in galls grown under N limitation. However, we investigated, if *dur3* might affect compatibility in –N conditions, but pathogenicity of the *dur3*-triple knockout strain was similar to wild type in both nitrogen regimes (Figure S5).

### Nit2 affects amino acid metabolism of galls during biotrophy

Although the impact of Nit2 deletion on pathogenicity was largely absent in both tested N regimes, effects on steady state levels of nitrate (see above) and some major amino acids in galls were evident, especially in the –N regime. First of all, this observation shows that Nit2-dependent changes in fungal metabolism affect overall gall metabolism, and secondly, this indicates that gall metabolism is affected stronger by Nit2 regulated genes in nitrogen deplete compared to nitrogen replete host plants. Significant differences between Δ*nit2* galls and wild type galls were observed for the nitrogen rich amino acids Asn and Gln, as well as for GABA. Asn and Gln are known to accumulate strongly in galls (Horst et al., 2010a) and represent major amino acids for source-sink transport in the phloem sap of maize (Wiener et al., 1991). We have demonstrated before that leaf galls represent strong sinks for organic nitrogen exported from lower systemic leaves (Horst et al., 2010a), so we can assume that Asn and Gln are unloaded together with phloem resident nitrate into the galls, where they are preferentially taken up by fungal hyphae. This idea is corroborated by the observation that after feeding of ^1^H-Asn, metabolization into organic acids and phosphorylated intermediates occurred much faster in galls compared to healthy control leaves (Horst et al., 2010b). Such assumptions should be interpreted with care, since we cannot discriminate the contribution of host and pathogen to the total free amino acid pool in galls. Since fungal colonization and gall size were comparable between the sampled Δ*nit2* and wild type galls, we can assume that the contribution of host metabolism is similar in both situations. Therefore, the specific reduction in steady state contents of Gln and especially Asn in Δ*nit2* galls compared to wild type galls in -N conditions strongly suggests that this reflects increased uptake of Asn by Δ*nit2* hyphae, probably as a compensation of the deficiency in nitrate uptake. Gln might even be converted to GABA in the apoplast and taken up by fungal hyphae as GABA, similar to the situation in *C. fulvum* discussed above (Solomon and Oliver, 2001; Solomon and Oliver, 2002). In support of this idea, the *U. maydis* homolog of the yeast GABA permease UGA4 (*um03522*) is one of the amino acid transport genes that show clear Nit2-dependent induction in –N during biotrophy, and *um03522* was also induced 1.8-fold in wild type hyphae in response to nitrogen depletion (Table S4). If we assume that GABA uptake is partially mediated by Nit2 in –N conditions, it cannot be explained, however, why GABA accumulation in Δ*nit2* galls is reduced compared to wild type galls under these conditions.

The final question is, whether we have obtained any indication that Δ*nit2* is nitrogen limited in -N conditions. First of all and most importantly, host colonization did not differ between the fungal genotypes in nitrogen deplete conditions, which speaks against nutrient limitation of the Δ*nit2* strain. In addition, steady state analysis failed to reveal differences in protein content between Δ*nit2* and wild type galls. Hence, it seems unlikely that availability of amino acid building blocks is more limited in Δ*nit2* than in the wild type.

But how can we explain the more than two-fold decrease in total protein content in gall tissue compared to mock leaves? In general, leaf protein is dominated by the photosynthetic apparatus. In galls however, the expression of photosynthetic genes is strongly repressed at 8 dpi (Döhlemann et al., 2008), and hence the accumulation of proteins involved in the photosynthetic machinery and the Calvin-Benson cycle will be strongly reduced. Likewise, the strong decrease in total protein content in mock leaves under nitrogen limitation compared to 1N conditions is likely to reflect reduced availability of amino acid building blocks for the synthesis of the photosynthetic machinery and enzymes of major carbohydrate metabolism.

## Conclusion

The role of potential nitrogen limitation for pathogenicity of fungal phytopathogens, especially for biotrophs, has long been debated in the last decades. Although this work does not provide final answers, whether nitrogen limitation occurs during the initial post-penetration phase and if this might affect virulence, we report several seminal findings in that

1. *Ustilago maydis* is unlikely to suffer nitrogen limitation in natural conditions
2. The ultilization of nitrate is dispensable for *U. maydis* biotrophy
3. The GATA transcription factor Nit2 regulates a different, but partially overlapping set of genes during biotrophy compared to saprotrophic sporidia
4. Nit2 and its effect on gene regulation during biotrophy is dispensable for full virulence
5. The utilization of nitrate and amino acids during biotrophy might be regulated based on availability

It remains a challenge for the future to determine synergistic and antagonistic effects of N limitation and *U. maydis* galls on metabolic reprogramming of maize leaves in order to increase our knowledge on how the fungal biotrophs subvert host metabolism. It seems especially rewarding to investigate how the flux of amino acids from host source leaves to strong N sinks like galls is affected in low N regimes. Likewise, resolving the contribution of (individual) amino acid, oligopeptide and nucleotide transporters to organic N supply to the fungus will improve our mechanistic understanding of the underlying metabolic reprogramming.

## Acknowledgements

Lars M. Voll, Christin Schulz and Alicia Fischer were funded by the German Research Council (Deutsche Forschungsgemeinschaft, DFG) via project VO 985/7-1. The authors are grateful for additional funding to Christin Schulz by short term scholarships from the Microcosm Earth Center (Marburg). Experimental support by Philipp L. Lopinski to Christin Schulz was funded by the Gender Equality Office of the Philipps-Universität Marburg.

The authors would like to thank Regine Kahmann (Max-Planck Institute for Terrestrial Microbiology) for the generous provision of *dur3-1,2,3* triple knockout strains. Critical reading of the manuscript and valuable suggestions by Johannes Freitag (University of Marburg) is also gratefully acknowledged. The authors would like to further acknowledge Christiane Rohrbach and Julian Rotfuß (University of Marburg) for skilled technical assistance and Özge Özbay (University of Marburg) for support in typesetting the manuscript.

## Author contributions

PLL, CS, AF, NR and NB performed the research and analyzed the data, TE and NB developed the HPLC method and analyzed the HPLC data, LMV conceptualized and designed the research, analyzed the data and wrote the paper with valuable input from TE.

## Supporting Information

**Supplemental Figure S1.** Validation of the FB1Δ*nit2* and FB2Δ*nit2* knockout by PCR on genomic DNA of single spore isolates.

**Supplemental Figure S2.** Nit2-dependent transcript accumulation of *U. maydis* genes in galls during biotrophy at 8 dpi.

**Supplemental Figure S3.** GO (gene ontology) term enrichment analysis for transcripts downregulated in Δ*nit2* compared to wild type in medium sized galls at 8dpi.

**Supplemental Figure S4.** Relative transcript abundance of all annotated *U. maydis* amino acid transporter genes in RNA-Seq data from medium size galls at 8 dpi.

**Supplemental Figure S5.** Pathogenicity of the *dur3-1,2,3* triple knockout mutant.

**Supplemental Table S1.** Composition of Hoagland Nutrient Solutions used to generate the three fertilization regimes 3N, 1N and –N.

**Supplemental Table S2.** Utilization of nitrogen sources by FB1Δ*nit2* and FB2Δ*nit2* sporidia in minimal medium.

**Supplemental Table S3.** Verification of transcript accumulation of selected Nit2 regulated genes in sporidia by qRT-PCR.

**Supplemental Table S4.** Downregulated *U. maydis* genes in medium sized Δ*nit2* versus wild type galls at 8 dpi, as determined by RNA-Seq. N=2

