## Supporting Material for "The *Ustilago maydis* transcription factor Nit2 regulates nitrate utilization during biotrophy and influences organic nitrogen metabolism in infected maize leaves under nitrogen limitation"

**Supporting Information**

**Table S1. Composition of Hoagland Nutrient Solutions** used to generate the three fertilization regimes 3N, 1N and –N (left to right). Plants were watered with the solutions 2-3 times per week, according to their demand.

| Component | Stock Solution | mL Stock Solution/1L | | |
|---|---|---|---|---|
| | | **3N** | **1N** | **-N** |
| 2M $KNO_3$ | 202g/L | 7.5 | 2.5 | 0 |
| 2M $Ca(NO_3)_2 \times 4H_2O$ | 236g/0.5L | 7.5 | 2.5 | 0 |
| Iron (Sprint 138 iron chelate) | 15g/L | 1.5 | 1.5 | 1.5 |
| 2M $MgSO_4 \times 7H_2O$ | 493g/L | 1 | 1 | 1 |
| 1M $NH_4NO_3$ | 80g/L | 3 | 1 | 0 |
| 1M $KH_2PO_4$ (pH 6.0) | 136g/L | 0.5 | 0.5 | 0.5 |
| 2M $CaCl_2$ | | 0 | 0 | 2.5 |
| Minors: | | 1 | 1 | 1 |
| $H_3BO_3$ | 2.86g/L | | | |
| $MnCl_2 \times 4H_2O$ | 1.81g/L | | | |
| $ZnSO_4 \times 7H_2O$ | 0.22g/L | | | |
| $CuSO_4$ | 0.051g/L | | | |
| $Na_2MoO_4 \times 2H_2O$ | 0.12g/L | | | |

**Table S2. Utilization of nitrogen sources by FB1Δ*nit2* and FB2Δ*nit2* sporidia in minimal medium.**

Growth of FB1, FB1Δ*nit2*, FB2, and FB2Δ*nit2* sporidia on minimal medium plates supplemented with selected single nitrogen sources. Sporidia dilution series of each genotype were plated and growth properties were rated 1 day after plating as normal growth (+), reduced growth (+-), strongly reduced growth (-) and no growth (--).

| | **FB1** | **FB1Δ*nit2*** | **FB2** | **FB2Δ*nit2*** |
|---|---|---|---|---|
| **ammonium** | + | + | + | + |
| **nitrate** | + | - | + | -- |
| **Ala** | + | - | + | -- |
| **Arg** | + | + | + | + |
| **Gln** | + | + | + | + |
| **Glu** | + | + | + | + |
| **Gly** | + | - | + | -- |
| **Leu** | + | - | + | -- |
| **Phe** | + | -- | + | -- |
| **Ser** | + | +- | + | - |

**Table S3. Verification of transcript accumulation of selected Nit2 regulated genes in sporidia by qRT-PCR.** FB1, FB2, FB1Δ*nit2* and FB2Δ*nit2* sporidia were transferred to nitrogen starvation (-N) or ammonium minimal medium (AMM) and harvested 2 h after transfer. Shown is the absolute fold change and standard error of –N compared to AMM control determined from two experimental replicates with three technical replicates. The normalized transcript amount of the indicates genes with *UmGAPDH* as a reference gene was determined with the primer pairs described by Horst et al. (2012).

| GeneID | Annotation | -N vs. AMM | | -N vs. AMM | |
|---|---|---|---|---|---|
| | | FB1 | FB2 | FB1Δ*nit2* | FB1Δ*nit2* |
| *um01756* | purine transporter | 1.96 ± 0.64 | 5.24 ± 1.71 | 0.58 ± 0.03 | 1.05 ± 0.03 |
| *um04577* | urea permease *dur3* | 19 ± 3 | 944 ± 306 | 4 ± 0.4 | 2 ± 0.1 |
| *um05889* | *ump2* | 1088 ±136 | 718 ± 125 | 64 ± 1.4 | 47 ± 1.4 |

**Table S4. Downregulated *U. maydis* genes in medium sized Δ*nit2* versus wild type galls at 8 dpi, as determined by RNA-Seq.** N=2

Medium sized galls of comparable size were harvested at 8 dpi for RNA-Seq analysis. See attached Excel file for data.

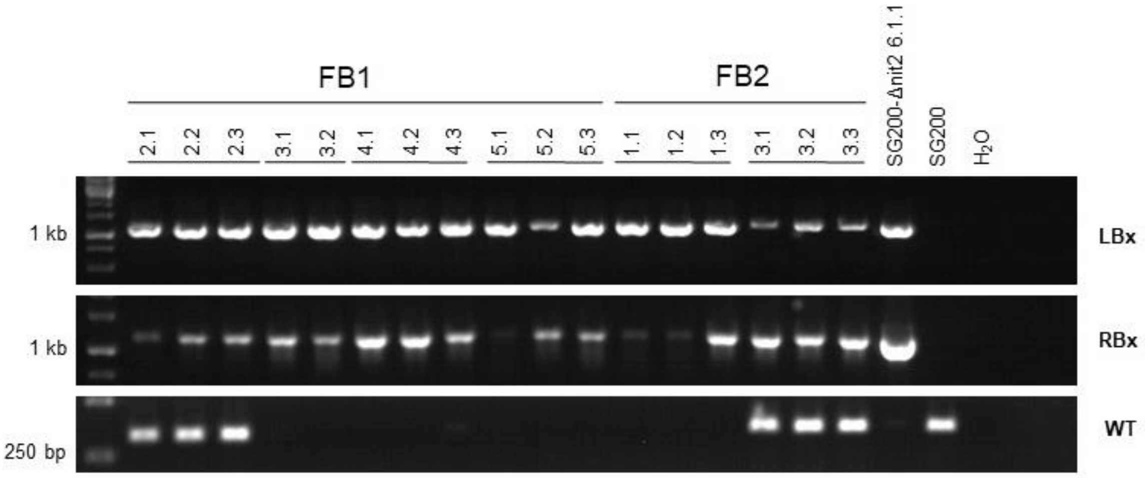


**Figure S1.** Validation of the FB1Δ*nit2* and FB2Δ*nit2* knockout by PCR on genomic DNA of single spore isolates.

The panels on top indicate strain (FB1 or FB2) and number of the single cell isolates tested, including one SG200Δ*nit2* and SG200 control on the right end. The left flank of the k.o. cassette insertion (LBx, top panel), right flank of the k.o. cassette insertion (RBx, middle panel) and the WT allele were detected by primer pairs nit2LBx 5'-CGAGTCTTTTCAGTCTTGTCTTTC-3' and hhn5.2 5'-CCGATGCAAAGTGCCGATAAAC-3' (for LBx) nit2RBx 5'-GGAGTGTCACAATTTCGGCTG-3' and hhn3.2 5'-GCTCAACTTTCATCGTGCCCAG-3' (for RBx) and wild type Nit2WT-fw 5'- GCCTCTCGAAGAAACAGTGG-3' and Nit2WT-rv 5'- AGAACGGGATCGACTGACAC-3' for the WT product.

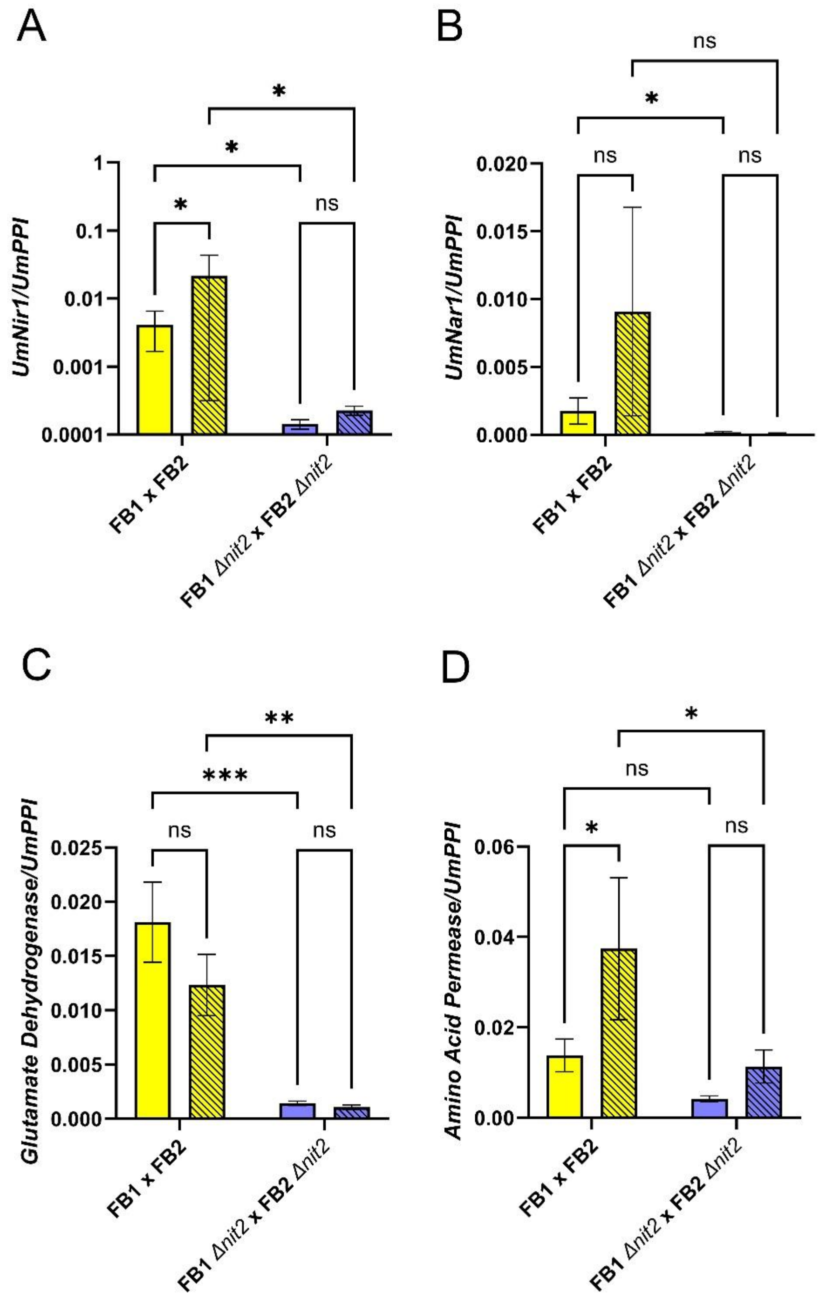


**Figure S2.** Nit2-dependent transcript accumulation of *U. maydis* genes in galls during biotrophy at 8 dpi.

Medium galls of comparable size were harvested at 8 dpi and transcript amounts for the *U. maydis genes* **(A)** nitrite reductase *nir1* (*um11104*), **(B)** nitrate reductase *nar1* (*um03847*), **(C)** NADP glutamate dehydrogenase (*um02801*) and **(D)** an amino acid permease (*um00056*) were quantified in a qRT-PCR relative to the *PPI* reference gene. Similar results were obtained when *PPI* was used as a reference gene (Figure S2). Values are means of 3-5 biological replicates ± SE. Statistical analysis was conducted with a two-way ANOVA a Fisher LSD post hoc test (*P < 0.05; **P < 0.01; ***P < 0.001; ****P < 0.0001).

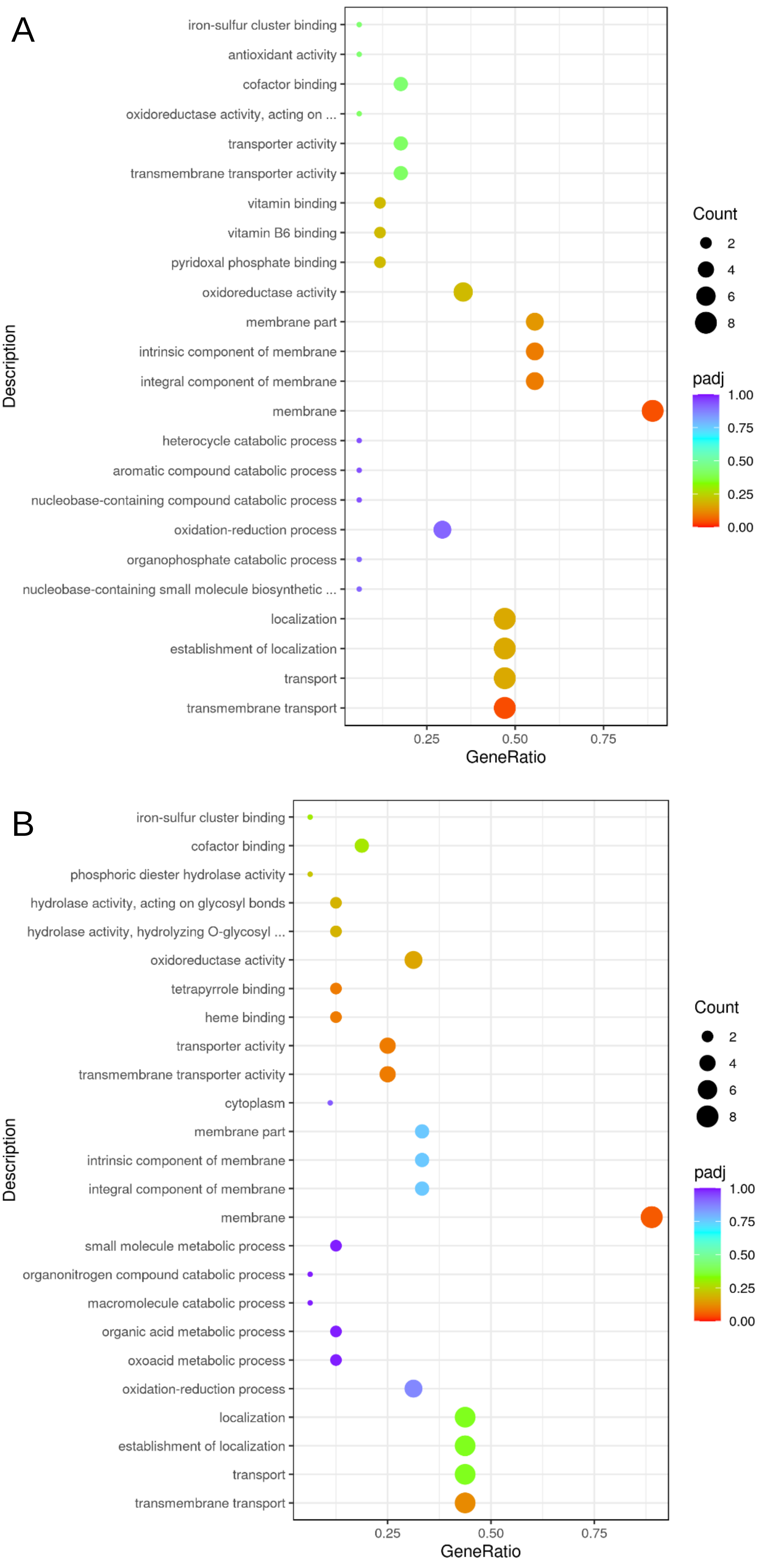
A
iron-sulfur cluster binding
antioxidant activity
cofactor binding
oxidoreductase activity, acting on ...
transporter activity
transmembrane transporter activity
vitamin binding
vitamin B6 binding
pyridoxal phosphate binding
oxidoreductase activity
membrane part
intrinsic component of membrane
integral component of membrane
membrane
heterocycle catabolic process
aromatic compound catabolic process
nucleobase-containing compound catabolic process
oxidation-reduction process
organophosphate catabolic process
nucleobase-containing small molecule biosynthetic ...
localization
establishment of localization
transport
transmembrane transport
Description
0.25
0.50
0.75
GeneRatio
Count
2
4
6
8
padj
1.00
0.75
0.50
0.25
0.00
B
iron-sulfur cluster binding
cofactor binding
phosphoric diester hydrolase activity
hydrolase activity, acting on glycosyl bonds
hydrolase activity, hydrolyzing O-glycosyl ...
oxidoreductase activity
tetrapyrrole binding
heme binding
transporter activity
transmembrane transporter activity
cytoplasm
membrane part
intrinsic component of membrane
integral component of membrane
membrane
small molecule metabolic process
organonitrogen compound catabolic process
macromolecule catabolic process
organic acid metabolic process
oxoacid metabolic process
oxidation-reduction process
localization
establishment of localization
transport
transmembrane transport
Description
0.25
0.50
0.75
GeneRatio
Count
2
4
6
8
padj
1.00
0.75
0.50
0.25
0.00

**Figure S3.** GO (gene ontology) term enrichment analysis for transcripts downregulated in Δ*nit2* compared to wild type in medium sized galls at 8dpi.

A – GO term enrichment among genes downregulated in Δ*nit2* versus wild type in 1N conditions. B – GO term enrichment among genes downregulated in Δ*nit2* versus wild type in -N conditions. Legends for gene count (bubble size) and adjusted p-value (color code) are shown next to the panels. On the x-axis, the relative fraction of downregulated relative to all genes in the respective GO term category is plotted.

Data are derived from the same experiment as the data shown in Table S3.

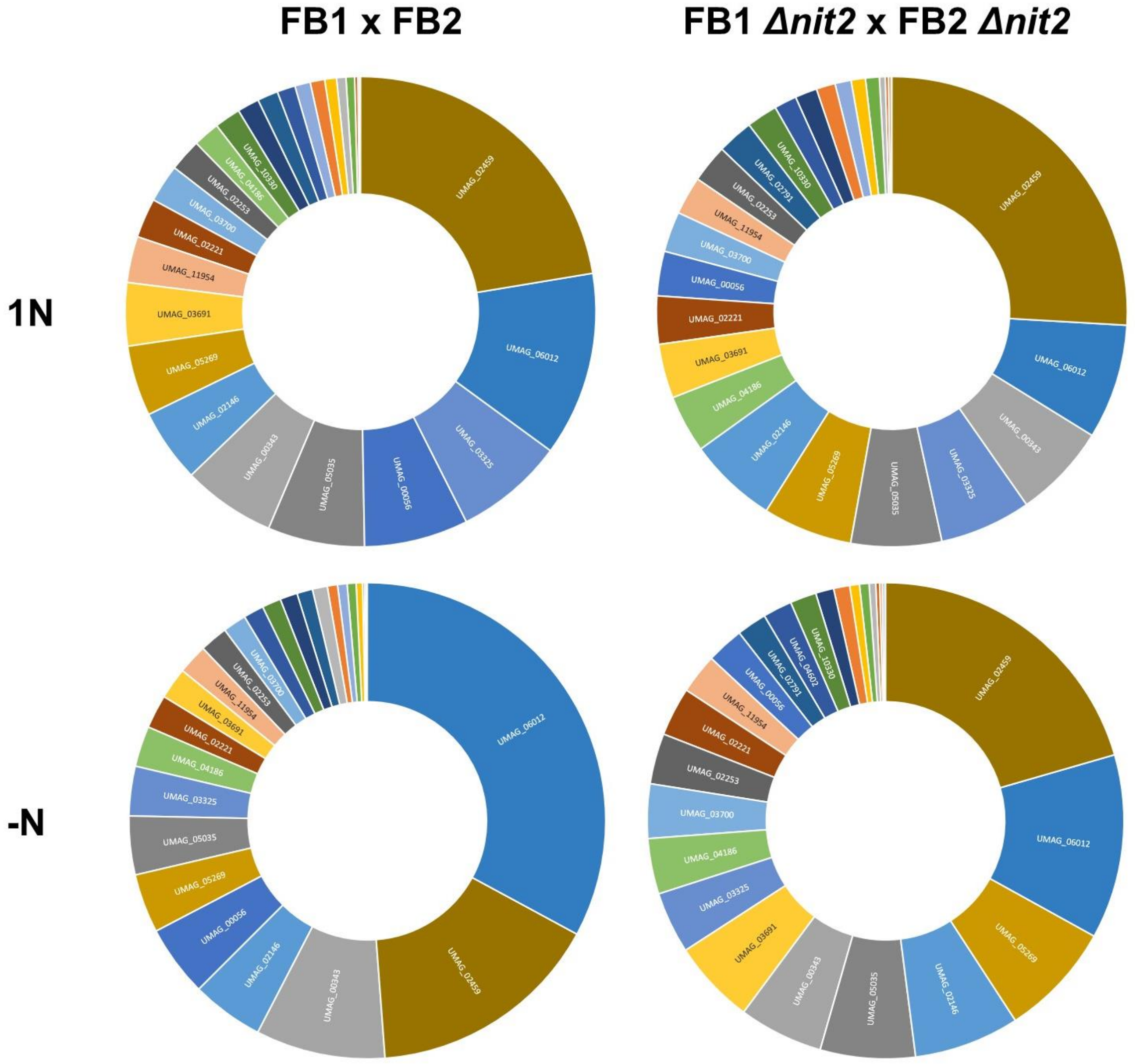


**Figure S4.** Relative transcript abundance of all annotated *U. maydis* amino acid transporter genes in RNA-Seq data from medium size galls at 8 dpi.

Mean FPKM values (N=2) were computed for all annotated amino acid transporter genes and the transcript amount of every individual gene relative to the total FPKM of all amino acid transporter genes is represented as pie charts. UMAG numbers are given for amino acid transporters with more than 1.5% share. 1N (upper row), -N (lower row), FB1 x FB2 (left column), FB1Δ*nit2* and FB2Δ*nit2* (right column). Data are derived from the same experiment as the data shown in Table S3.

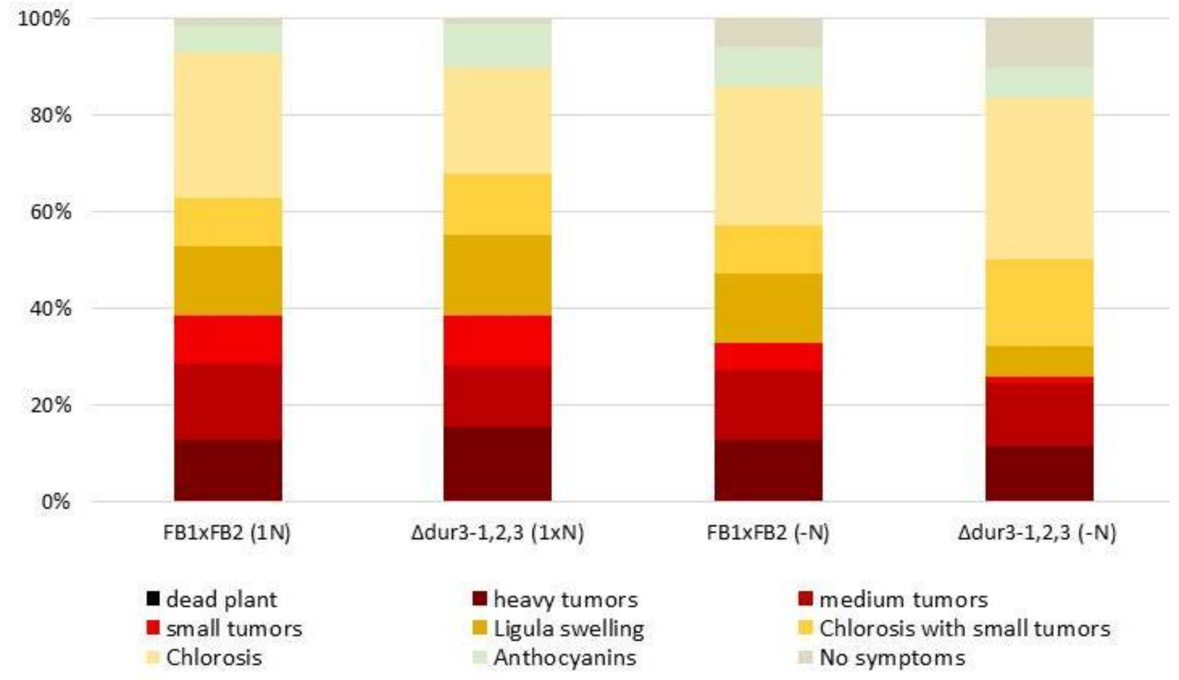


**Supplemental Figure S5.** Disease index of the *dur3-1,2,3* triple knockout mutant at 8 days post infection dpi. To this end, 7 days old plants had been infected with a mixture of the indicated sporidia at a titer of OD=1 each. N= 70-78.
